# HT-SIP: A semi-automated Stable Isotope Probing pipeline identifies interactions in the hyphosphere of arbuscular mycorrhizal fungi

**DOI:** 10.1101/2022.07.01.498377

**Authors:** Erin E. Nuccio, Steven J. Blazewicz, Marissa Lafler, Ashley N. Campbell, Anne Kakouridis, Jeffrey A. Kimbrel, Jessica Wollard, Dariia Vyshenska, Robert Riley, Andy Tomatsu, Rachel Hestrin, Rex R. Malmstrom, Mary Firestone, Jennifer Pett-Ridge

## Abstract

**Background:** Linking the identity of wild microbes with their ecophysiological traits and environmental functions is a key ambition for microbial ecologists. Of many techniques that strive to meet this goal, Stable Isotope Probing—SIP—remains the most comprehensive for studying whole microbial communities *in situ*. In DNA-SIP, active microorganisms that take up an isotopically heavy substrate build heavier DNA, which can be partitioned by density into multiple fractions and sequenced. However, SIP is relatively low throughput and requires significant hands-on labor. We designed and tested a semi-automated DNA-SIP pipeline to support well-replicated, temporally-resolved amplicon or metagenomics experiments that enable studies of dynamic microbial communities over space and time. To test this pipeline, we assembled SIP-metagenome assembled genomes (MAGs) from the hyphosphere zone surrounding arbuscular mycorrhizal fungi (AMF), in combination with a ^13^CO_2_ plant labelling study.

**Results:** Our semi-automated pipeline for DNA fractionation, cleanup, and nucleic acid quantification of SIP density gradients requires six times less hands-on labor compared to manual SIP and allows 16 samples to be processed simultaneously. Automated density fractionation increased the reproducibility of SIP gradients and reduced variation compared to manual fractionation, and we show adding a non-ionic detergent to the gradient buffer improved SIP DNA recovery. We then tested this pipeline on samples from a highly-constrained soil microhabitat with significant ecological importance, the AMF fungal hyphosphere. Processing via our quantitative SIP pipeline confirmed the AMF *Rhizophagus intraradices* and its associated microbiome were highly ^13^C enriched, even though the soils’ overall enrichment was only 1.8 atom% ^13^C. We assembled 212 ^13^C-enriched hyphosphere MAGs, and the hyphosphere taxa that assimilated the most AMF-derived ^13^C (range 10-33 atom%) were from the phlya Myxococcota, Fibrobacterota, Verrucomicrobiota, and the ammonia oxidizing archaeon genus *Nitrososphaeara*.

**Conclusions:** Our semi-automated SIP approach decreases operator time and errors and improves reproducibility by targeting the most labor-intensive steps of SIP—fraction collection and cleanup. Here, we illustrate this approach in a unique and understudied soil microhabitat—generating MAGs of active microbes living in the AMF hyphosphere (without plant roots). Their phylogenetic composition and gene content suggest predation, decomposition, and ammonia oxidation may be key processes in hyphosphere nutrient cycling.

## BACKGROUND

Stable isotope probing ‘SIP’ approaches, where active microbes are identified via incorporation of stable isotopes (^13^C, ^15^N, ^18^O) into their biomarkers, cells, DNA or RNA, are among the most powerful methods in microbial ecology since they can identify the most relevant active microbes and their specific ecophysiology traits in natural, ‘wild’ settings [1–7]. Broadly speaking, SIP refers to any technique where microorganisms that have consumed substrates enriched in rare stable isotopes (e.g. ^13^C, ^15^N, ^18^O) are identified based on the resulting isotopic enrichment of their nucleic acids, proteins, and metabolites (reviewed in [5, 8]). However, DNA-SIP—where isotopically enriched DNA is separated from unenriched nucleic acids via isopycnic separation in cesium chloride—is the most commonly used SIP approach, typically in conjunction with 16S rRNA gene or shotgun metagenome analysis. A cornerstone of many seminal studies of microbial biogeochemical cycling, DNA-SIP has been used to identify populations involved in decomposer food webs [9], consume plant root exudates [10, 11], degrade pollutants [12] and C1 compounds [13], oxidize ammonia [14], fix N2 [15, 16], or to characterize population growth, survival and mortality in mixed communities [17, 18].

Quantitative stable isotope probing (qSIP) and techniques such as ‘high resolution’ SIP (HR-SIP) [19] are expansions of the original SIP concept that combine density gradient ultracentrifugation with mathematical models designed to improve the quantification (sensitivity and specificity) of isotope enrichment [6, 20]. When used with a ^18^O-enriched ‘heavy water’ addition, qSIP enables calculation of growth and mortality rates of individual taxa, since cells and viruses incorporate oxygen from water during nucleic acid synthesis, quantitatively reflecting cell division (DNA synthesis) and metabolism (RNA synthesis) [21–23]. Recent qSIP studies have used the method to illustrate how wild microbial communities are shaped by evolutionary history [24, 25], soil temperature and warming [26, 27], amendments of water and nutrients [18, 28] and trophic relationships amongst bacterial predators and their prey [29].

While the majority of SIP and qSIP studies have focused on 16S rRNA gene profiles, targeting active populations with shotgun sequencing (metagenomes and metatranscriptomes) provides greater opportunity for inference of genomic potential and whole genome-scale data analysis [30–32], define microbial guilds [33], and provide insights into cross-kingdom interactions (including virus-host matching) [23, 34]. But SIP-metagenomics is a daunting prospect for many research groups, as multiple metagenome datasets are generated from each initial microbiome DNA sample, creating financial and computational limitations that constrain experimental scope. Sieradzki et al. (2020) [35] explain the relationships between sample replication, enrichment and the precision achievable with a given number of SIP density fractions (and suggest 9 fractions is typically ideal). Additionally, processing SIP density gradients is a relatively low throughput process and requires significant hands-on labor, making it onerous to conduct the well-replicated, temporally resolved experiments needed to study dynamic microbial community activities over space and time. Historically, SIP studies have used few replicates due to the laborious nature of the technique. To address this short-coming, we have designed a high-throughput SIP (HT-SIP) pipeline for processing SIP density gradients, which automates fractionation, partially automates fraction cleanup, and automates the preparation of samples for nucleic acid quantification. We have now tested replicates of this pipeline at both LLNL and the JGI for over two years, running 1000s of samples.

To validate and demonstrate the utility of our HT-SIP pipeline on an important yet challenging sample set, we targeted the ‘hyphosphere’ soil microhabitat using ^13^C labeling—the area under direct influence of arbuscular mycorrhizal fungal (AMF) hyphae. Arbuscular mycorrhizal fungi (members of the Glomeromycota) form obligate symbiotic associations with 80% of all land plants [36], and in exchange for plant carbon (C), supply their host with essential nutrients such as N and P [37, 38] and water [39]. Intriguingly, AMF are capable of stimulating decomposition of soil organic matter (SOM) and dead plant material [40–42], but do not have the enzymatic repertoire to decompose SOM themselves. As such, the importance of interactions with the soil microbiome is potentially critical [43], and previous research suggests that AMF modify their surrounding soil litter-decomposing microbial community in order to acquire nitrogen derived from SOM, and transport it to the host plant [38, 44]. However, these interactions occur at such a small spatial scale (hyphae are ca. 1.5 – 18 µm in diameter [45]) that they are extremely difficult to measure and monitor. Using SIP, in conjunction with ^13^CO_2_ labeling of a plant host inoculated with AMF, we tracked plant-fixed carbon through AMF hyphae and into the surrounding hyphosphere microbiome.

To generate SIP-metagenomes from the AMF-hyphosphere, we ^13^CO_2_ labeled the wild annual grass, *Avena barbata*, inoculated with the AMF *Rhizophagus intraradices* in sterile sand. The microcosms contained a separate hyphal chamber with live soil that only AMF hyphae could access, from which hyphal aggregates were collected and extracted for ^13^C-hyphosphere SIP processing. We used our semi-automated pipeline to process samples from this microhabitat, and produced high quality libraries and MAGs even while using an unusually low starting DNA input for SIP separations (350 ng DNA). Our work demonstrates that automation not only saves operator time and improves reproducibility of SIP processing, but is also suitable for analysis of low DNA quantities and downstream amplicon and metagenomics analysis. The ^13^C-hyphosphere MAGs assembled in this study are a key advance for dissecting trophic interactions in the AMF hyphosphere.

## METHODS

### Density Gradient Separations

HT-SIP validation experiments were conducted using 1-5 µg DNA for SIP density gradient separations (below, amounts and DNA sources are specified per experiment). To separate DNA based on isotopic enrichment, DNA was added to 150 µL 1xTE buffer mixed with 1.00 mL gradient buffer, and 4.60 mL CsCl stock (1.885 g mL^-1^) with a final density of 1.725-1.730 g mL^-1^. Samples were loaded into ultracentrifuge tubes (5.1 mL, Quick-Seal Round-Top Polypropylene Tube, Beckman Coulter) and spun at 20°C for 108 hours at 176,284 RCFavg (equivalent to 176,284 x *g*) in a Beckman Coulter Optima XE-90 ultracentrifuge using a VTi65.2 rotor, following a previously described protocol [17, 35] to create density gradients.

### High-Throughput SIP (HT-SIP) Pipeline

To automate the labor-intensive steps of SIP—density gradient fractionation, cleanup, and quantification—we combined a series of robotic instruments. Following cesium chloride (CsCl) density gradient separation in an ultracentrifuge, we automated fractionation by connecting an Agilent Technologies 1260 Isocratic Pump and 1260 Fraction Collector to a Beckman Coulter Fraction Recovery System (see Supplemental Figure S1 for schematic and parts list). In this system, each sample is separated into 22 fractions (∼236 µL each). CsCl is displaced in the ultracentrifuge tube by pumping sterile water at 0.25 mL min^-1^ through a 25G needle inserted into the top of the ultracentrifuge tube, and the sample fraction exits via a side port needle inserted into the bottom of the tube. We maintain pressure between 1-1.8 bar; pressures above this indicate the system is clogged. The gradient medium fractions are dispensed into 96-well deep well plates (2 ml square well plates with v-bottoms, Stellar Scientific) by the Agilent Fraction Collector. Four SIP tubes are fractionated into a single deep-well plate (88 wells) and the final row is left empty for PicoGreen quantification standards. At the beginning of the day and after every four gradients, we clean the fractionation tubing with water using a “wash spacer” to bypass the fraction recovery system (see Supplemental Figure S1). The density of each fraction is measured manually using a Reichart AR200 digital refractometer fitted with a prism covering to facilitate measurement from 5 µL, as previously described [46].

DNA in each density fraction is then purified (desalted) and concentrated using a Hamilton Microlab STAR liquid handling robot, which we have programmed to automate PEG precipitations using a previously published protocol [47], with modifications for 96-well plates. We configured our robot deck to process four plates; this allows a maximum of 16 SIP samples to be processed simultaneously (4 samples per plate). Following fractionation, the robot adds 2 volumes of 30% PEG 6000 (in 1.6 M NaCl) and 35ul of 1:5 diluted Glycoblue (Invitrogen, Thermo Fisher) to each well. Plates are then manually sealed and mixed thoroughly by vortexing and manual shaking, pulsed down briefly, and incubated at room temperature in the dark overnight. To precipitate the DNA, we spin the four plates at 4198 RCF for 5 hours at 20°C in an Eppendorf 5920R centrifuge using a S-4xUniversal-Large rotor. The plates are then placed back in the Hamilton robot, which removes the PEG by pipetting and rinses the pellets using 950ul 70% ethanol. Plates are manually sealed, gently mixed by vortexing, and centrifuged at 4198 RCF for 1.5 hours at 20°C to stabilize the DNA pellets. We aspirate the ethanol with the robot, and then manually place the plates upside down on a paper towel to drain remaining ethanol. The plates are then returned to the robot to dry for 15 minutes, whereafter the robot automatically resuspends the DNA pellets in 40 µL of 1x Tris-EDTA (pH 7.5); 10 mM Tris-HCl may be used for applications sensitive to EDTA. Finally, plates are manually sealed and stored at -20°C.

Finally, DNA concentration of each fraction is quantified with a PicoGreen fluorescence assay (Invitrogen, Thermo Fisher). Picogreen quantification plates are prepared in triplicate on a Hamilton Microlab STAR robot, where each plate contains a row for the standard curve. Samples are mixed with the PicoGreen reagent in a 96-well intermediate mixing plate, and then distributed into three 96-well PCR plates for fluorescence analysis. Plate fluorescence is measured in a CFX Connect Real-Time PCR Detection System (Bio-Rad), and the fluorescence values for the three technical replicate plates are averaged to determine DNA concentration.

### Validation of HT-SIP using Manual SIP

To validate the automated steps of our HT-SIP pipeline, we compared fractionation and PEG precipitations using both manual and automated methods. Automated fractionation was performed as described above, and manual fractionation was conducted with a Beckman Coulter fraction recovery system as previously described [17]. Samples were fractionated into approximately 22 fractions, although the number of fractions recovered by manual SIP typically varies despite identical run conditions.

To compare automated verses manual PEG precipitations, 4 ug soil DNA (extracted from a sample collected at the Hopland Research and Extension Center in Hopland, CA 38°59’35"N, 123°4’3"W) was added per density gradient. For manual samples, following manual fractionation, PEG precipitations were conducted in microcentrifuge tubes as previously described [47] using published centrifuge speeds and times, which we note are faster than those used for our HT-SIP plate-based method.

### Increasing DNA-SIP recovery using non-ionic detergents

Absorption of DNA to polypropylene tubes can lead to substantial sample loss, especially for DNA in high ionic strength solutions [48], but this concern can be mitigated by adding non-ionic detergents [48]. Since the ultracentrifuge tubes used in DNA-SIP protocols are made of polypropylene and CsCl is a high ionic strength solution, we tested whether adding the non-ionic detergents Tween-20 and Triton-X to density gradient buffer improved DNA recovery. To identify the optimal concentration of detergent for DNA-SIP recovery, we tested additions in the range of 0.0001 – 1% for Tween-20 and 0.0001 – 0.1% for Triton-X and compared DNA recovery versus the standard density gradient formulation. 1 ug of *E. coli* genomic DNA (Thermo Scientific) was added to density gradients and processed using the HT-SIP pipeline (n=3 gradients per condition). After identifying that adding 0.0001% Tween-20 had the highest percent DNA recovery, we assessed 0.0001% Tween-20 additions to a larger set of soil DNA samples (101 SIP tubes total) using our HT-SIP pipeline. We added 4 µg of soil DNA (from Hopland, CA soil) to these gradients.

### Validation of HT-SIP Pipeline: Hyphosphere ^13^CO_2_ Labeling and Harvest

AMF hyphosphere soil was ^13^C-labeled in ^13^CO_2_ plant growth chambers; details on the microcosm design and growth conditions are documented in [39]. Briefly, *Avena barbata* seedlings were planted in the ‘plant compartment’ of two-compartment microcosms and grown for 10 weeks. The plant compartment was separated from the ‘no-plant compartment’ by a 3.2 mm air gap to prevent root exudates or dissolved organic C from travelling via mass flow between compartments. Both sides of the air gap had nylon mesh that either allowed hyphae but excluded roots (18 μm mesh), or that excluded both hyphae and roots (0.45 μm mesh).

In the plant compartment, a sterile sand mix (1:1 volumes of sand and clay, plus 78 mg of autoclaved bonemeal) was inoculated with 26 g of whole inoculum of *Rhizophagus intraradices* (accession number AZ243, International Culture Collection of (Vesicular) Arbuscular Mycorrhizal Fungi (INVAM), West Virginia University, Morgantown, WV). The no-plant compartment contained a mixture of 1:1 volumes of live soil (from Hopland CA) and sand plus 78 mg of autoclaved bone meal.

The microcosms were incubated in growth chambers in the Environmental Plant Isotope Chamber (EPIC) facility, located in the Oxford Tract Greenhouse at UC Berkeley, in temperature-controlled growth chambers with a multiplexed ^13^CO_2_ delivery system monitored by IRGA and Picarro CO_2_ analyzers. For this study, three microcosms with 18 μm mesh (^13^C AMF permitted in the no-plant compartment, termed ‘^13^C-hyphosphere’) and three microcosms with 0.45 μm mesh (^13^C AMF excluded from the no-plant compartment, termed ‘^13^C no-AMF control’, for IRMS analysis only) were continuously ^13^CO_2_-labeled for 6 weeks during weeks 5-10. Six additional microcosms remained in a natural abundance CO_2_ atmosphere for the full ten weeks; of these, the three ^12^C microcosms with 18 μm mesh (^12^C AMF permitted in the no-plant compartment, termed ‘^12^C-hyphosphere’) served as the ^12^C-hyphosphere SIP controls, and three ^12^C microcosms with 0.45 μm mesh (^12^C AMF excluded from the no-plant compartment) were for IRMS analysis only. AMF-specific Sanger sequencing of the plant compartment (roots, sand) as well as the air-gap indicated the planted compartment only contained the initial mycorrhizal inoculum [39].

At the beginning of week 11, all microcosms were destructively sampled. Soil was placed in Whirl-Pak bags, flash frozen in liquid nitrogen, and stored at -80°C. To collect hyphosphere microbial communities, visible hyphal aggregates with hyphosphere soil attached were collected from the no-plant compartments using tweezers under a dissecting microscope for +AMF microcosms. For no-AMF controls, the soil mix was sampled the same way, except no hyphae were visible. During the microcosm harvest, plant shoots, roots, and soil from the planted and unplanted chambers were placed in paper envelopes and then dried 60°C for 72 hours for ^13^C IRMS analysis. These samples were finely ground, weighed, and analyzed for total C and ^13^C abundance by dry combustion on a PDZ Europa ANCA-GSL elemental analyzer interfaced to a PDZ Europa 20-20 isotope ratio mass spectrometer (Sercon Ltd., Cheshire, UK). Stated precision by the manufacturer for ^13^C is 0.1 per mil.

### Hyphosphere HT-SIP Density Gradient Separations

DNA was extracted from 250 mg of hyphal aggregates using the DNeasy PowerSoil kit (Qiagen). We added 350 ng of ^13^C- and ^12^C-hyphosphere hyphosphere DNA (n = 3 each) to density gradients and ultracentrifuged as described above, and then fractionated, precipitated, and quantified using the HT-SIP pipeline. DNA pellets were resuspended in 10mM Tris-HCl (pH 7.5) because the low DNA mass in each fraction required us to use a large fraction volume during sequence library creation and would have resulted in higher than recommended EDTA concentrations ( < 0.1 mM EDTA final concentration).

### Metagenomic Sequencing Library Preparation and Sequencing

Since the overall soil ^13^C-enrichment was relatively low in these samples, we sequenced metagenomes from 14 fractions per sample to increase the chance of detecting taxa with significant ^13^C-enrichment; Sieradzki et al. [35] have shown that sensitivity is correlated with isotopic enrichment. Fractions with low concentrations of DNA at the beginning (fractions 3-6) and end of the gradient (fractions 19-21) were combined and concentrated prior to sequencing using an Amicon Ultra-0.5 30KDa filter (Millipore Sigma) [49]; fractions 7-18 were sequenced without concentration. Whole genome metagenomic libraries were prepared at LLNL using the Illumina DNA Flex library kit (now called Illumina DNA Prep, Illumina Inc., Santa Clara, CA) using 1 ng of sample DNA and 12 cycles of amplification. The libraries were dual indexed with Illumina Nextera DNA CD indexes following the manufacturer’s recommended protocol and quantified using a Qubit broad range dsDNA assay (Thermo Fisher Scientific, Waltham, MA). Library insert sizes were determined via Agilent Tapestation with the D5000 High Sensitivity assay (Agilent Technologies, Santa Clara, CA). Equimolar amounts of each library were pooled together. The pooled libraries’ sizes and concentrations were verified using the D5000 High Sensitivity assay (Agilent Technologies, Santa Clara, CA).

The library was diluted and denatured as described [50] to a final concentration of 375 pM (2% phiX and 98% library). The library pools were sequenced with NextSeq 1000/2000 P2 or P3 reagents (Illumina Inc., Santa Clara, CA) as paired end 2 X 150 cycles on an Illumina NextSeq2000 Sequencer at Lawrence Livermore National Laboratory. In total, 2.6x10^9^ read pairs passed quality filtering, with a mean of 3.1x10^7^ read pairs per sample/fraction and a range of 3.0x10^6^ to 1.1x10^8^ read pairs.

### Metagenome Assembly

Our prior research indicates that coassemblies of all SIP gradient fractions from biological replicates yield the best metagenome-assembled genomes (MAGs) [31, 51]. Metagenomic sequences were loaded into IMG v5.1.1. We created individual assemblies using metaspades [52] and a single co-assembly including all fractions using MetaHipMer v2 [51]. MetaHipMer produced more contigs (Supplemental Table S2), so we proceeded with this assembly. Sequences were mapped to the assembly using bbmap [53], binned with Metabat v2.12.1, and then bins were refined with metaWRAP v1.2.1 [54]. MAG quality was determined by CheckM v1.1.3 [55], taxonomy determined by GTDB-tk v1.50 with version r202 of the GTDB taxonomy database [56] and final metrics are reported using Minimum Information about a Metagenome Amplified Genome (MIMAG) standards [57]. MAG abundances were determined using bbsplit [53]. The sequenced genome of *Rhizophagus irregularis* DAOM 197198 was included in the bbsplit reference library to map AMF sequences (RefSeq assembly accession: GCF_000439145.1) [58].

The genomic capacities of high- and medium-quality MAGs of highly ^13^C enriched hyphosphere taxa were determined by annotating into functional categories with Patric subsystems [59] and KEGG orthologs from KofamKOALA [60]. Carbohydrate active enzyme (CAZyme) gene homologs were determined using dbCAN HMM analysis [61]. Protease homologs were determined using blastp against the MEROPS v.12.1 database [62]; an initial screen was performed using the “merops_scan.lib” database, and positive hits were then compared to the larger “pepunit.lib” database, which is a non-redundant library of all peptidase units and inhibitor units in MEROPS.

### Quantitative SIP Analysis

We applied quantitative SIP (qSIP) calculations to estimate the atom percent excess (APE) ^13^C enrichment for each taxon following procedures detailed in Hungate et al. [20] with adjustments for metagenome-assembled genomes instead of 16S rRNA genes [31, 34]. qSIP uses the number of reads per fraction across the SIP gradient to estimate the APE of each taxon within the enriched sample compared to the natural abundance control. To estimate the amount of taxon-specific DNA per fraction, we calculated the proportion of the library attributable to the MAG and multiplied by the DNA content per fraction ([MAG_counts / total_library_counts] * ng_fraction_DNA). Since our reference database of MAGs is incomplete, and does not include genomes for eukaryotes and fungi other than the AMF *R. intraradices*, we normalized MAG counts to total sequencing library counts because this includes the portion of the library with no sequenced representatives. After the MAG sequence counts were converted into estimates of DNA mass (ng), weighted average densities (WAD) for each MAG were calculated for each gradient. The differences in WAD between the ^13^C and ^12^C gradients were used to estimate the atom percent excess for each MAG [20]. All MAGs were present in all three replicates. To be considered ^13^C-enriched, the 90% lower confidence interval of a MAG’s median APE-^13^C needed to be >0% APE-^13^C.

## RESULTS

### Reproducibility of automated fractionation

Automated fractionation increased the reproducibility of fractionation; the buoyant densities of automated samples had a more consistent linear fit across gradients (r^2^ = 0.984, n = 24 gradients) compared to manually fractionated gradients (r^2^ = 0.908, n = 22 gradients) (**Figure 1**). With manual fractionation, the sample to sample variability of fraction density increased in fractions collected later in the process (less dense) and resulted in a more variable number of fractions even though identical fractionation conditions were used (range = 15 – 27 fractions); in contrast, automated fractionation consistently resulted in 22 fractions.

**Figure 1.**
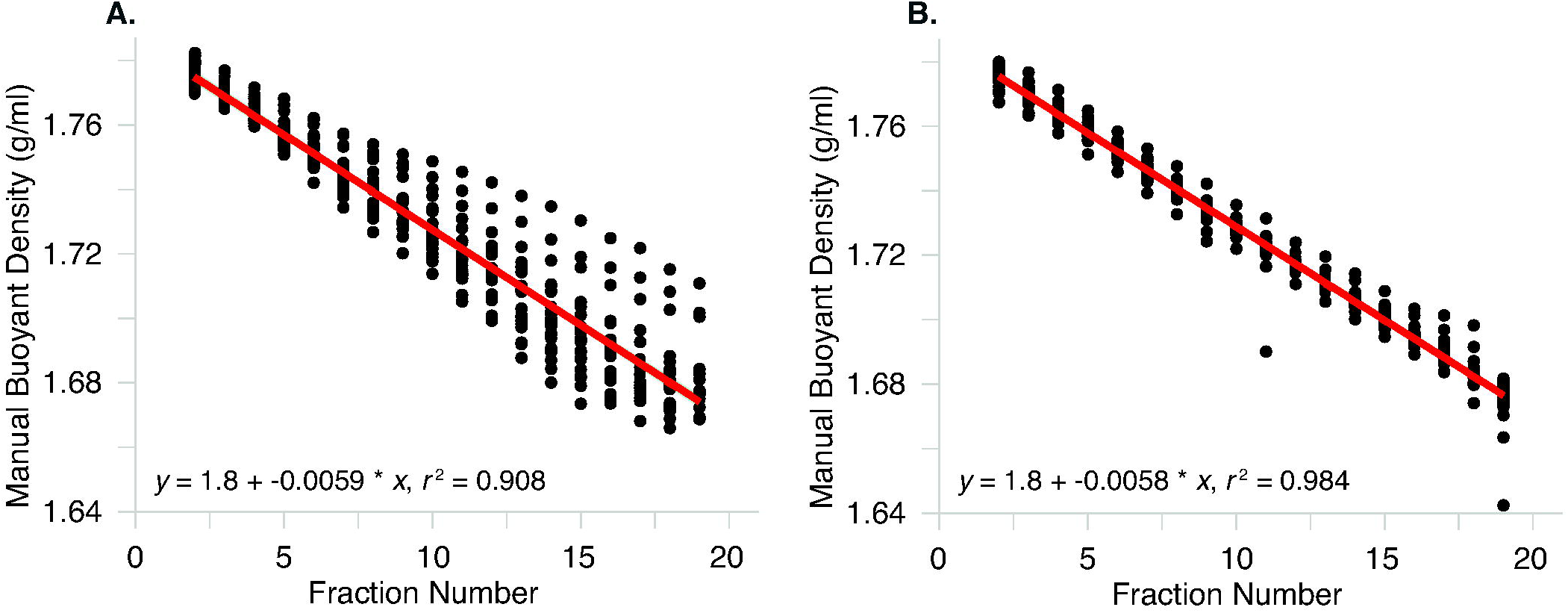
Comparison of manual versus automated fractionation of SIP density gradients. Fractionation is the process of dividing a SIP density gradient into multiple fractions, where the initial fractions at the bottom of the tube are the heaviest and the final fractions at the top of the tube are the lightest. Buoyant density (g/ml) for each fraction is measured via refractometry and is represented by a single dot. **(A)** For “manual” fractionation, 22 independent density gradients were fractionated by visually counting and collecting droplets in microcentrifuge tubes using the method described in Blazewicz et al [17]. The number of fractions collected from manual density gradient was variable (range 15-27 fractions per gradient); only fractions 2-19 are displayed. **(B)** For automated fractionation, 24 independent density gradients were fractionated robotically using an Agilent Technologies fraction collector, which automatically divides the gradients into fractions of a set volume (∼236 µl) and dispenses them into a 96-well plate. Automated density gradients consistently produced 22 fractions per gradient. Fractions at the beginning and end of the gradient (fractions 1, 20-22) were excluded as these densities are altered by the water used to displace the gradient and are not typically used for molecular analysis.

### Effect of non-ionic detergents on automated DNA recovery

Post-fractionation DNA recovery is a key concern for DNA-SIP studies, especially for samples where total DNA volume is limited. In our initial testing of >1000 environmental DNA samples (at LLNL and JGI), we noticed DNA percent recovery tended to improve when larger masses of DNA were added to the density gradient (3-5 µg) and hypothesized that DNA was being lost by an adsorption mechanism. We therefore tested how adding the non-ionic detergents Tween-20 and Triton-X affected DNA recovery in density gradients, which can mitigate potential DNA loss by adsorption to polypropylene tube walls [48]. Adding Tween-20 at final concentrations ≤ 0.01% reliably increased the recovery of 1 µg pure culture *E. coli* DNA, and Tween-20 at 0.0001% yielded the highest overall DNA recovery (84 ± 5%) compared to a no-detergent control (64 ± 5% recovery) (Supplemental Table S1). Triton-X only improved DNA recovery at a final concentration of 0.0001% (74 ± 3%).

**Table 1.** Time comparison for manual versus automated SIP fractionation, cleanup, and DNA quantification. Estimates are based on processing 22 fractions per SIP tube. The “Hands-on” columns indicate the time an individual must actively manipulate the samples, while “total” columns indicate the total time required for the entire process. Manual cleanup and quantification time estimates are based on processing a single SIP gradient and assume maximum processing speed. Automated fraction cleanup time is based on the time required to fractionate a single tube using the Agilent Infinity Fraction Collector. Automated sample cleanup and DNA quantification times are based on the processing times for the Hamilton STAR liquid handling robots. The † symbol indicates that 16 SIP gradients are processed simultaneously in 4 plates (4 gradients per plate), and the “per 1 sample” times are calculated by dividing the total time by 16. “Batch” processing times are the times required to process 16 density gradients.

| SIP Processing Steps | Manual Time (hours) |  | Automated Time (hours) |  |
| --- | --- | --- | --- | --- |
|  | Hands-on | Total | Hands-on | Total |
| Fraction Collection | 1 | 1 | 0.2 | 0.6 |
| Sample Cleanup | 0.4 | 1 | 0.04† | 0.5† |
| DNA Quantification | 0.5 | 0.6 | 0.06† | 0.13† |
| Total Time (per 1 sample) | 1.9 | 2.6 | 0.3† | 1.2† |
| **Batch Time (per 16 samples)** | **30** | **42** | **4.8** | **20** |

Using soil DNA in the absence of Tween-20, we had higher DNA recovery with manual processing (measured after the precipitation step) relative to the automated protocol (**Figure 2a**). When 0.0001% Tween-20 was included in the density gradient buffer, soil DNA recovery with the automated protocol was comparable to the manual approach (**Figure 2**). For the automated protocol, adding Tween-20 significantly increased soil DNA yields by a factor of 1.8 from 35 ± 4% to 64 ± 3% (mean ± 95% CI; p< 0.01 for one-tailed t-test) (**Figure 2b**).

**Figure 2.**
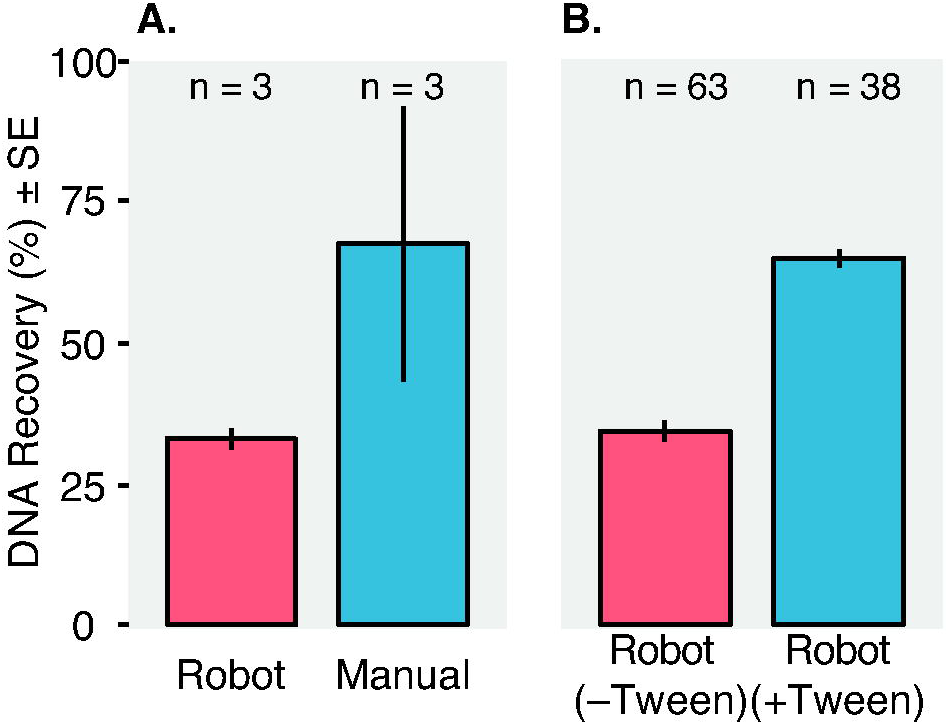
DNA recovery comparison for manual and semi-automated PEG precipitation methods, and the impact of adding a non-ionic detergent (Tween-20) to the SIP gradient buffer. After a density gradient is fractionated, each fraction needs to be desalted prior to quantification and sequencing analysis, which can be accomplished using nucleic acid precipitations. (A) We compared “manual” PEG precipitations (n=3 SIP gradients), where each fraction is precipitated in microcentrifuge tubes by an individual (as per Blazewicz et al. [17]), and semi-automated or “robot” PEG precipitations (n=3 SIP gradients), where a Hamilton STAR liquid handling robot performs the precipitations in 96-well plates. This process is semi-automated because some steps require assistance from an individual (e.g., transferring plates to a centrifuge for DNA sedimentation). (B) We tested how adding Tween-20 to the density gradient mixture impacts DNA recovery for a large SIP experiment analyzing soil DNA; all samples were processed semi-automatically by the robot. Tween-20 was added to a subset of the samples (+Tween, n = 38 SIP gradients) or processed using our standard density gradient buffer without Tween-20 (–Tween, n = 63 SIP gradients). Both experiments were conducted using 4 µg soil DNA per SIP gradient; recovery was calculated by summing recovered DNA (measured by Picogreen) in the recovered density fractions post-cleanup and dividing by the initial DNA input. Error bars represent the standard error of the mean.

### SIP automation time savings

In HT-SIP, automating multiple steps in sample SIP processing substantially increased the overall speed of the procedure and decreased the amount of manual or “hands-on” labor required. Using automation during fraction collection, sample cleanup, and DNA quantification, it is manageable to process 16 samples in parallel (this batch number is constrained by the typical number of spaces available in an ultracentrifuge rotor). Overall, HT-SIP requires half as much technician time to fully process 16 samples (42 h versus 20 h for 16 samples), and 6.25 times fewer hands-on hours compared to serially processing single samples manually (**Table 1**)); within a work week, this represents 3.2 days of hands-on labor saved. Manual estimates assume no parallelization and are based on the minimum amount of time required to process a single SIP gradient under ideal conditions in our laboratory.

### HT-SIP validation: Hyphosphere soil

To demonstrate the HT-SIP pipeline on an important yet challenging sample set (low DNA, low overall enrichment), we targeted hyphosphere soil (the area under direct influence of arbuscular mycorrhizal fungal (AMF) hyphae) using ^13^C labeling of the annual grass *Avena barbata* in a two-compartment microcosm. After six weeks of labeling, roots in the planted compartment of ^13^C-AMF microcosms were highly enriched (41.3 ± 1.9 atom% ^13^C). Root biomass in the ^12^C microcosms was unenriched (1.1 ± 0.001 atom% ^13^C). In the no-plant compartment of ^13^C-AMF microcosms, bulk analysis of the soil showed it was slightly enriched (1.8 ± 0.1% atom% ^13^C), whereas in the no-AMF control microcosms (where AMF were excluded from the no-plant compartment), the soil ^13^C content was not statistically different from the ^12^C control soil (1.1 ± 0.001% versus 1.1 ± 0.0003%, respectively, *p* < 0.001). This indicates that ^13^CO_2_ diffusion into the soil from the labeling chamber headspace was minimal, there was no significant autotrophic ^13^C-fixation in the no-plant compartment, and that enriched ^13^C detected in the hyphosphere was transferred primarily by the AMF hyphae.

To identify ^13^C enriched metagenome assembled genomes (MAGs) in the hyphosphere, we used the HT-SIP pipeline on DNA extracted from soil-covered hyphal aggregates (termed, “hyphosphere”) collected from the no-plant compartment in both ^13^C- and ^12^C-AMF microcosms. SIP density separations of total DNA showed evidence of only slight ^13^C-enrichment, as seen by the small increase in weighted average density (WAD) between ^12^C and ^13^C samples (**Figure 3a**). After mapping the SIP metagenomic reads to the *R. irregularis* genome, we calculated atom percent excess (APE) using qSIP, and found the AMF DNA was significantly ^13^C-enriched (23 atom% ^13^C) (**Figure 3b**).

**Figure 3:**
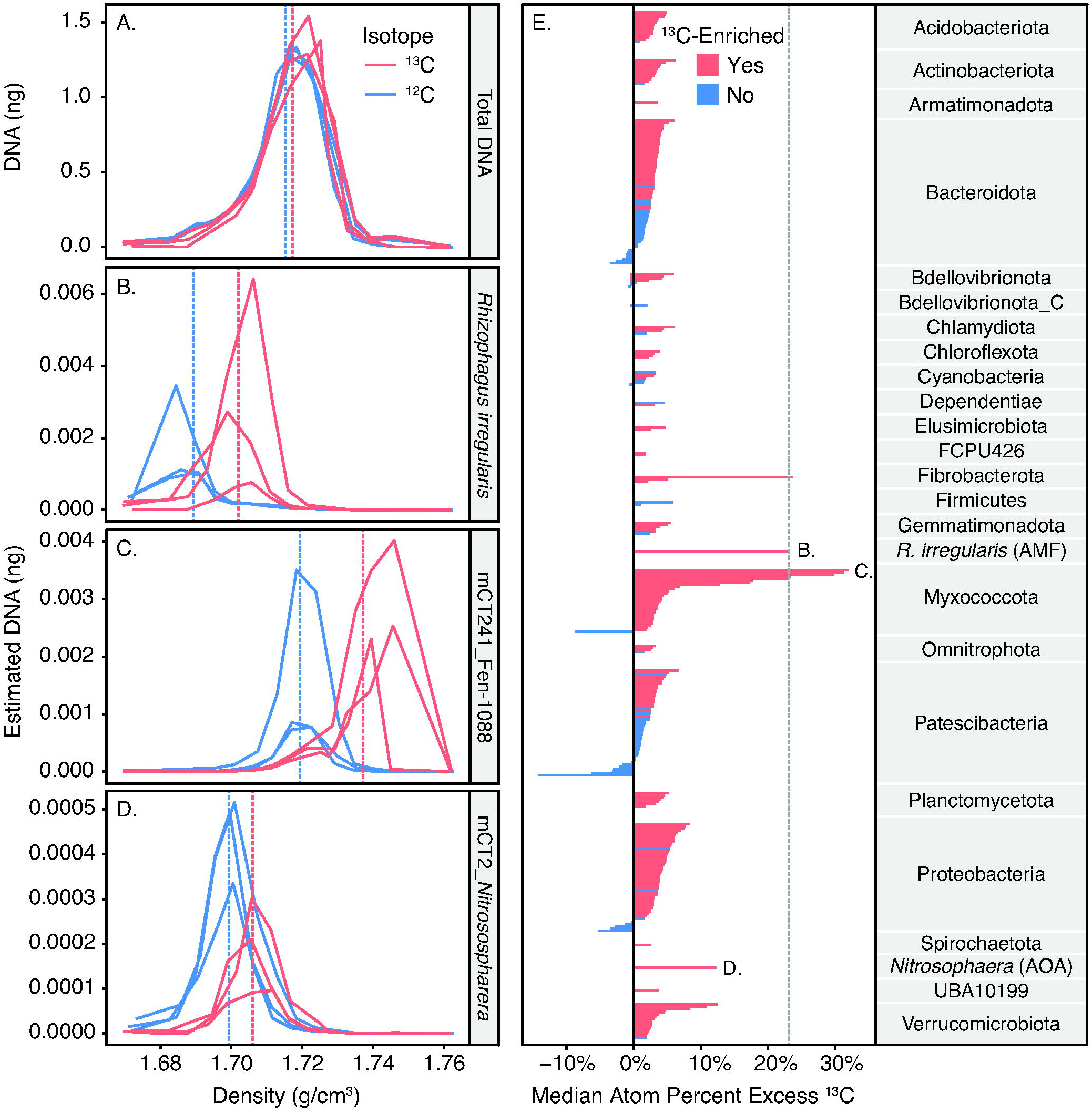
^13^C-Hyphosphere Metagenome Assembled Genomes (MAGs) isolated by SIP-metagenomics. Concentration versus fraction density of **(A)** total DNA extracted and SIP fractionated from a ^13^C-hyphosphere soil (red lines) and ^12^C-hyphosphere control soil (blue lines); n=3. Dashed lines are the weighted average density (WAD) of the DNA of the replicate gradients for each isotope. Within this gradient, we used qSIP to estimate taxon-specific DNA masses for **(B)** the AMF host *Rhizophagus intraradices*, |and two MAGs recovered from the AMF hyphosphere: **(C)** a Myxococcota MAG (mCT241_Fen-1088) (the most ^13^C-enriched organism detected), and **(D)** an archaeal ammonia oxidizer (mCT2_*Nitrososphaera*). Taxon-specific DNA masses were estimated by multiplying a fraction’s total DNA mass by taxon relative abundance (e.g., MAG counts divided by metagenomic library counts). **(E)** Estimated median atom percent excess (APE) of all assembled MAGs, which were calculated based on the difference in weighted average density between ^13^C-hyphosphere samples and ^12^C-hyphosphere control samples. Red bars indicate the 212 MAGs that had significantly greater ^13^C enrichment than 0 (lower 90% CI bound greater than 0), and blue bars indicate the MAGs that were unenriched (lower 90% CI bound below 0). The dashed gray line indicates the APE of the AMF, *Rhizophagus intraradices*, which supplied ^13^C to the hyphosphere chamber. Taxa are grouped by phylum, and letters indicate the APE of the taxa shown in panels B-D.

### Hyphosphere-qSIP metagenome assembly and binning

All fractions were co-assembled using MetaHipMer2, which produced 14.4 assembled Gbp > 1 kb and 1.6 assembled Gbp > 10 kb. Compared to single fraction assembly using metaspades, MetaHipMer2 produced 3.3 and 3.2 times more assembled Gbp > 1 kb and Gbp > 10 kb, respectively (Supplemental Table S2). We therefore proceeded with the MetaHipMer2 assembly (which did not require bin dereplication). Overall, 71.2 % of the sequence reads mapped to the MetaHipMer2 assembly. Binning produced 299 medium- and high-quality MAGs; completeness, percent contamination, and MAG genome size are available in Supplemental Table S3. Three MAGs were 100% complete, including taxa in the orders Cytophagales (mCT1; Bacteroidota), Nitrososphaerales (mCT2; Archaea), and Pedosphaerales (mCT3; Verrucomicrobiota). The phyla with the most MAGs assembled were Bacteroidota (68), Proteobacteria (51), the candidate phylum radiation Patescibacteria superphylum (50), and Myxococcota (30). On average, 13.3 ± 7.6% SD of the sequence reads mapped to the MAGs.

### Genomic potential of ^13^C-enriched MAGs in the AMF hyphosphere

The soil microbial community in the ^13^C AMF hyphosphere was significantly isotopically enriched; of the 299 assembled MAGs, 212 were significantly ^13^C enriched, indicating they consumed plant-fixed ^13^C transported by the AMF hyphae (**Figure 3e**). Of these, 43 MAGs were moderately enriched (> 5-10% APE-^13^C) and 12 MAGs were highly enriched (>10% APE-^13^C), which included 8 Myxococcota MAGs, a Fibrobacterota from the family UBA11236 (mCT95), an ammonia oxidizing archaeon (AOA) from the family *Nitrososphaera* (mCT2), and two MAGs from the Verrucomicrobiota family Opitutaceae (mCT7, mCT160) (**Figure 4**).

**Figure 4:**
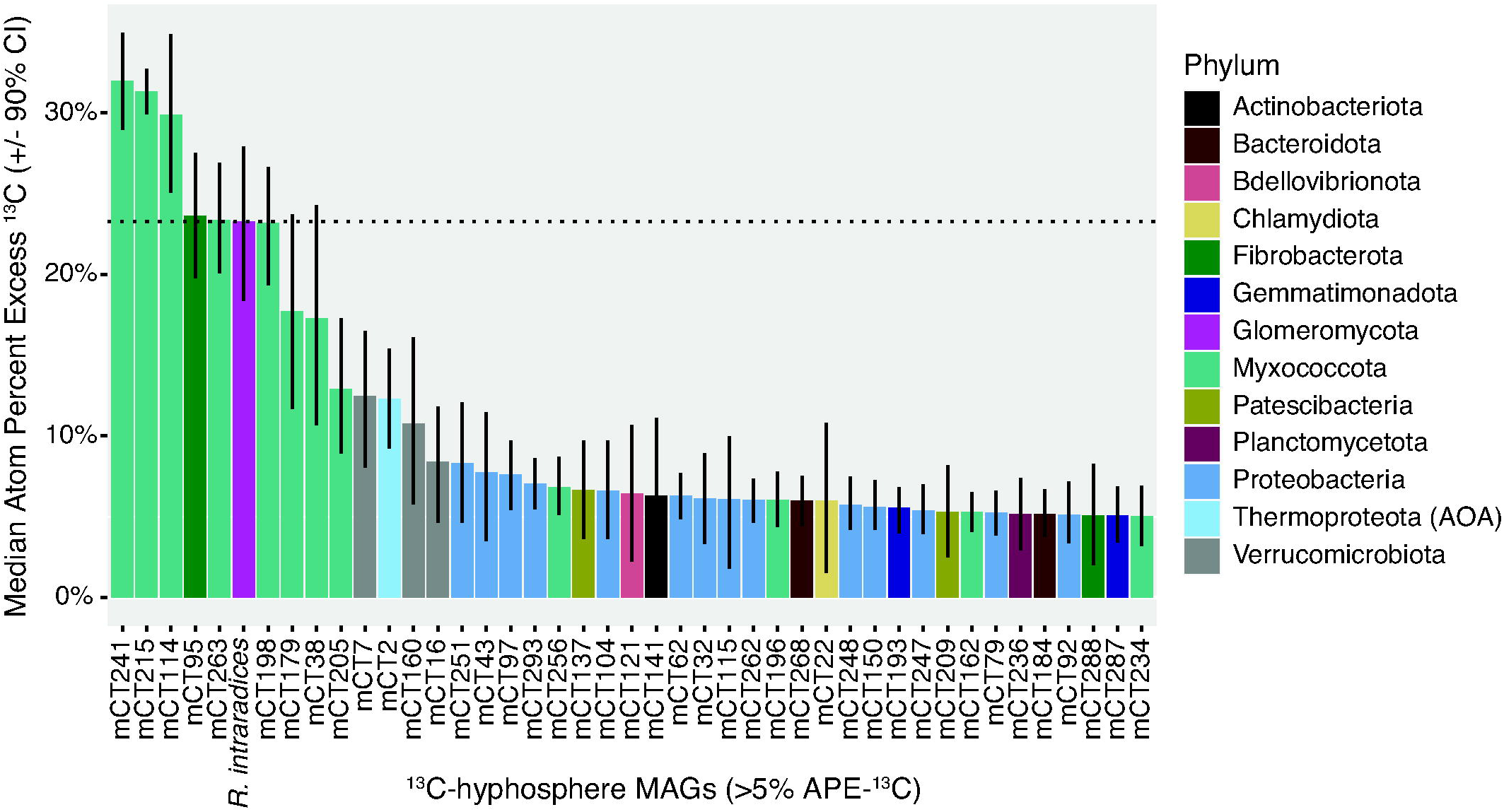
Ranked enrichment of highly enriched MAGs in the AMF ^13^C-hyphosphere, as estimated by qSIP. MAGs displayed have >5% atom percent excess (APE) ^13^C and are colored by phylum affiliation. Dashed line indicates the APE-^13^C of the AMF, *Rhizophagus intraradices*. Error bars represent the 90% confidence interval. A full list of MAGs and their isotopic enrichment is available in Supplemental Table 3.

We used comparative genomics of carbohydrate degradation genes (CAZymes), as well as other genes, to assess possible roles for these MAGs in the microbial food web based on their genomic potential. Except for the AOA, highly enriched MAGs had multiple homologs of PL6 genes (alginate and non-alginate polysaccharide lyases [57]) and GH109 genes (glycoprotein α-N-acetylgalactosaminidases) (Supplemental Table S4); PL6 family enzymes were disproportionately abundant in highly ^13^C-enriched MAGs (12 of 29 MAGs).

Myxococcota are facultative microbial predators that can subsist by predation or saprotrophy [63]. Eight Myxococcota MAGs ranged in APE-^13^C from 12-32%, and seven of these were from the little-known Polyangia family Fen-1088 that is known only from metagenomic sequencing (only 31 MAGs in the Genome Taxonomy Database (GTDB)) (accessed June 2022) [64]. Similar to many Myxococcota [65], Fen-1088 have high GC contents (68-71%) and large genome sizes (3.5-7.8 Mbp) (Supplemental Table S3). Three highly-enriched Fen-1088 MAGs (mCT241, mCT215, mCT114) were ca. 10 APE-^13^C more enriched than the AMF and ranged from 30-32% APE-^13^C (**Figure 3c**, mCT241). Compared to a set of Mycococcota type species genomes [65], these three Fen-1088 MAGs contain similar amounts of genes associated with predation, such as potential cell lysis CAZyme families (7 GH13 genes, 5-7 GH23 genes) and protease genes (203-225 MEROPS protease homologs) (Supplemental Table S5). Two of the MAGs contain a chitinase gene (mCT114, mCT215) (Supplemental Table S6). However, these highly enriched MAGs were enriched in glycoside hydrolases and polysaccharide lyases that are atypical for Myxococcota type species [65], such as GH29 (α-fucosidases), GH109 (α-N-acetylgalactosaminidases), GH8 (β-1,4 linkages, such as those found in plant cell walls), PL6 (alginate and non-alginate lyases [66]), and PL14 (lyases of unknown function, including poly(b-mannuronate) lyase). These genomic differences are also apparent across the Myxococcota MAGs from this study, where the highly ^13^C-enriched Fen-1088 MAGs consistently contained more PL6 and GH29 homologs than less-enriched Myxococcota (**Figure 5**).

**Figure 5:**
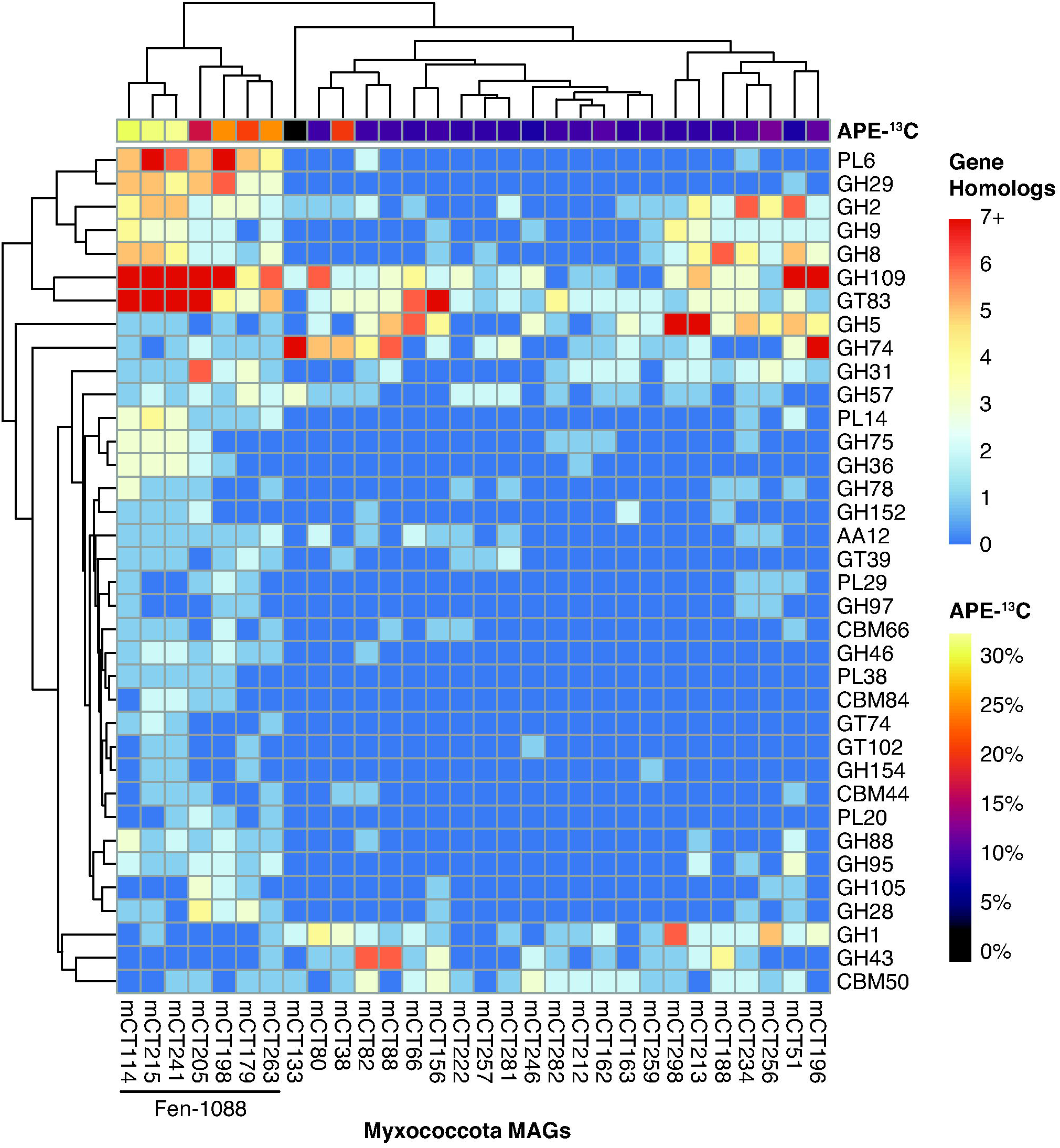
Genomic comparison of CAZy family homologs identified in Myxococcota MAGs assembled in this study. Rows indicate the number of gene homologs (red-blue color scale) detected per CAZy family, columns indicate the associated MAGs. Atom percent excess ^13^C (APE-^13^C) estimated by qSIP is presented in the top row (yellow-purple-black color scale). Columns and rows were clustered using one-dimensional hierarchical clustering based on genomic content. CAZy families displayed had significantly more or less genes detected in the Fen-1088 MAGs compared to the rest of the Myxococcota MAGs (student’s t-test, p <0.05); full results are available in Supplemental Table S6. Acronyms: GH = glycoside hydrolase; PL = polysaccharide lyase; CBM = carbohydrate binding module; GT = glycosyltransferase; AA = auxiliary activity.

Fibrobacterota mCT95 had the same APE as the AMF (23% APE-^13^C) and has chitinolytic potential (9 genes with GH18 domains and 2 genes with GH19 domains, 6 of which were annotated as chitinases (EC 3.2.1.14)). This Fibrobacterota MAG also contained GH74 genes for lysing β-1,4 glucan linkages in plant cell wall polysaccharides.

The AOA *Nitrososphaera* mCT2 (**Figure 3d**) was enriched at 12% APE-^13^C and contained genes for ammonia oxidation (amoABC) and nitrite reductase (nirK), as well as the marker gene for the 3-hydroxypropionate/4-hydroxybutyrate cycle for autotrophic C fixation (hydroxybuteryl-CoA dehydratase) [67]. The *Nitrososphaera* MAG had low glycoside hydrolase content (only 7 genes), no polysaccharide lyase genes, but many glycosyltransferase genes (24 genes).

Finally, two Opitutaceae MAGs were 11-12% APE-^13^C enriched. These genomes contained multiple genes with CAZy domains for processing glucose- and galactose-based uronic acids, such as genes for lysing β-1,4 linkages in polyglucuronic acid (PL20), hydrolizing α-1,4 glycosidic linkages in polygalacturonic acid (GH28), and hydrolyzing glucuronic and galacturonic acid monomers (GH105). Polygalacturonases lyse the pectin in plant cell walls [68], while polyglucuronases target the cell walls of bacteria, fungi, and algae [69]. The Opitutaceae MAGs also contained multiple copies of GH29 α-fucosidases. A detailed list of taxonomy and isotopic enrichment for all MAGs is available in Supplemental Table S3, and a comparison of the CAZyme gene content for MAGs with >10% APE-^13^C is available in Supplemental Table S4.

## DISCUSSION

Stable Isotope Probing is a powerful technique for resolving population demographics, functional traits, and ecological interactions of active microorganisms *in situ,* without the need for cultivation. However, SIP is not as broadly used as it could be, particularly for sequence-intensive metagenomics and transcriptomics studies, because it is relatively low throughput, time-consuming, and highly manual. Therefore, SIP has rarely been applied with well-replicated, temporally-resolved experimental designs necessary to reveal evolving microbial community activities over space and time. We have created a high throughput semi-automated SIP pipeline, ‘HT-SIP’, which enables substantially higher sample throughput, reduces labor costs, and makes it more feasible to invest effort in high-risk low-biomass samples that can be used to target the metagenomes of specific function-based subpopulations in complex microbial communities. Since establishing our pipeline (replicated at both LLNL and JGI), we have run more than 1000 samples from a diverse array of sites, including samples from boreal, temperate grassland, agricultural, and tropical forest habitats. This scalable pipeline automates density gradient fractionation, fraction cleanup, and quantification, and we show this method decreases operator time, reduces operator error, and improves reproducibility. Using HT-SIP, we were able to target an important microhabitat, the AMF hyphosphere, and examine potential trophic interactions in the fungal hyphosphere based on ^13^C-enrichment.

### Hyphosphere-SIP: Processing samples with low isotopic enrichment

Assessing a sample’s isotopic enrichment (e.g. via mass spectrometry), is often used as a pre-screen factor when deciding whether or not to proceed with SIP density gradient separations, and the number of replicates and SIP fractions to be sequenced. Samples with lower isotopic enrichment require more replicates and fractions to detect taxon-specific enrichment [35]. Based on our soil IRMS data and SIP data alone (**Figure 3a**), we typically would not have proceeded with the hyphosphere samples collected from our ^13^C AMF study, due to their low overall enrichment. However, since AMF are obligate biotrophs and the plant biomass was highly ^13^C enriched, we hypothesized that AMF hyphae and their surrounding hyphosphere might also be enriched, and that the low bulk ^13^C hyphosphere value might have been caused by dilution from a large ^12^C soil background, along with our inability to physically sample this microhabitat precisely. We thus proceeded with a DNA-SIP strategy of sequencing all fractions (14 total) to increase the chance of detecting enriched taxa. While initial IRMS data and total DNA-SIP separations suggested SIP might be impracticable on these samples due to insufficient ^13^C label, instead, taxon-specific qSIP indicated a highly enriched ^13^C-AMF signal.

### Dissecting trophic interactions in the AMF hyphosphere

Paradoxically, AMF are capable of stimulating decomposition of SOM and detritus [37, 38], but do not have the enzymatic repertoire to decompose SOM themselves. We and others have hypothesized that AMF collaborate with their soil microbiome to mineralize organic nutrients [43]; previous research shows the presence of AMF modifies the litter-decomposing microbiome [38] and the hyphosphere microbiome synergistically increases the amount of nitrogen AMF transfer back to the plant host [70]. The ^13^C-AMF hyphosphere MAGs we identified in this study are a key advance, enabling us to test hypotheses related to these phenomena and determine the molecular mechanisms that underpin them. Because SIP-metagenomes target DNA from active organisms that recently consumed an isotopically-labeled substrate, their genomic content represents the machinery that runs the microbial food web and provides a means to dissect potential trophic interactions.

### Predation in the hyphosphere

Using qSIP-estimated MAG atom percent excess, we examined potential trophic interactions in the fungal hyphosphere based on ^13^C-enrichment. Intriguingly, Myxococcota from the poorly-characterized family Fen-1088 were the most enriched taxa detected in the hyphosphere. While Myxococcota are a known component of AMF hyphosphere communities [71], their functional role has not been previously determined. The particularly high atom percent ^13^C enrichment of the Fen-1088 family suggests they are either directly feeding upon C exuded by the AMF, or are predators which target AMF hyphae or other microbes in the AMF hyphosphere. The Myxococcota phylum contains many facultative predators that can subsist by consuming microbial or plant organic matter [63], and have a broad prey range including bacteria and fungi [72, 73]. Since our Myxococcota MAGs have high GC content and large genomes, it is unlikely that they are AMF endosymbionts, which often have reduced genomes. Previous GWAS analysis indicates that Myxococcota have many prey-specific genes, rather than a general set of antimicrobial genes, which likely enable them to target a broad prey range [74]. Our analysis of the Fen-1088 MAGs also points to a large arsenal of proteases that may be used to consume prey, these are also found in many Myxococcota type species [65]. However, the relative lack of GH18 chitinase genes suggests this family may not be chitinolytic or may have low chitin degradation efficiency, as highly efficient chitinolytic organisms are thought to produce multiple types of chitinases [75, 76].

While the Fen-1088 MAGs are similar to other Myxococcota genomes, in that they contain a large array of CAZymes for carbohydrate and polysaccharide degradation, they also contain multiple copies of genes with CAZyme domains that are uncommon in Myxococcota type species. Most of these CAZymes are hypothetical proteins, but the presence of these CAZyme domains hints at potential function. We consistently found two enzyme groups, GH29 and PL6, in Fen-1088 MAGs. The GH29 group is known to contain alpha-fucosidases, which remove terminal L-fucoses from oligosaccharides or their conjugates[77]. Many biomolecules are fucosylated [78]— polysaccharides, glycoproteins, and glycolipids can have attached fucoses [77]. AMF hyphae are well-known for exuding glycoproteins and related compounds, that appear to play a key role in soil carbon stabilization. Fucose can be exuded and tightly attached to the AMF hyphal surface; in one previous study, fucose represented 3.5-5% of the mass of poly- and mono-saccharides that were exuded or could be gently hydrolyzed from the fungal surface [79]. *Rhizophagus irregularis* also exudes fucosylated lipo-chitooligosaccharides [80], which may be signaling molecules used when establishing a symbiosis with a plant host [81].

Much less is known about polyspecific enzyme family PL6; these enzymes are abundant in Fen-1088, and in all our MAGs with >10% APE-^13^C except the AOA MAG. PL6 contains alginate lyases and non-alginate lyases [66] that cleave ß(1-4) linkages within polysaccharides built from mannuronic and guluronic acids [66], such as alginate produced by brown algae and some bacteria, or between these acids and other building blocks for non-alginates. While these polysaccharides have been detected in soil, their origin is difficult to determine. Bacteria in the genera *Pseudomonas* and *Azotobacter* species can produce a bacterial form of alginate [82]. Little is known about the composition of mycorrhizal exo- and endo-polysaccharides, so it is not possible to rule out that these enzymes are acting on an AMF-produced polysaccharide. Mannose, the sugar from which mannuronic acid is built, is abundant in the mono- and polysaccharides bound to the surface of AMF hyphae [79], but we have no information about how these monosaccharides are polymerized. Further research on both the polysaccharides in the AMF hyphosphere and the feeding preferences of the Fen-1088 Myxococcota are needed to determine the nature of this relationship.

Stable isotope measurements can benefit food web studies, where the ^13^C enrichment of an organism is used as a conservative indicator of the ^13^C enrichment of the substrate they consume [29, 83]. In some cases, predators can be more enriched than their prey. For example, in a previous qSIP study, viral DNA was more ^13^C-enriched than the microbial host, likely because the host was using more recently assimilated C to support viral replication [34]. Similarly, in a recent qSIP meta-analysis, predators assimilated ^18^O or ^13^C at higher rates than non-predators [29]. We note that isotopic concentration by predators due to isotopic fractionation (ca. 1 per mil per trophic level for C [83]) is not relevant to our study, where we have used tracer levels of isotope enrichment. In our study, the Fen-1088 MAGs were more ^13^C enrichment than the AMF, even though AMF were the only source of ^13^C in the no-plant compartment. Myxococcota likely became more enriched than the mean AMF enrichment by consuming ^13^C-rich compounds recently transported from the plant host, such as newer hyphae or fresh exudates. At the time of harvest, plant root bulk enrichment was 41 atom% and AMF hyphae were 23 atom% (estimated by qSIP), but we expect that recently produced hyphae had a higher isotopic signature, more similar to the plant host (i.e. DNA extracted from all present AMF biomass would include hyphae produced earlier part of the ^13^CO_2_ labeling period when the plant was less ^13^C enriched).

### Interactions between AMF and ammonia oxidizing archaea (AOA)

The relatively high ^13^C enrichment of the *Nitrosophaera* MAG (12 APE ^13^C) suggests this AOA was intimately associated with the AMF, or nearby biota that obtained substantial quantities of ^13^C from the AMF (e.g., *Fibrobacterota, Opitituceae, Myxococcota*). While AOA are dominantly autotrophic, there is some evidence they can use organic C [84–86]. Because we did not detect ^13^C enrichment in soil from hyphae-free controls, it is unlikely non-specific subsurface diffusion of ^13^CO_2_ gas contributed to autotrophic ^13^C assimilation. Therefore, we conclude that AMF played a key role in the transport of plant photosynthate C to the AOA—either through direct (AMF è AOA) or indirect (AMF è other biota è AOA) pathways.

The nature of multitrophic interactions between AMF, AOA, and other soil biota remains largely unknown. AMF can induce changes in AOA community structure [87], with positive, negative, and negligible effects on AOA abundance [87–90]. Positive interactions between AMF and AOA may be driven by C supply or hyphosphere soil acidification. Low soil pH promotes AOA abundance and activity [84]. Negative interactions may be driven by competition for ammonia, which can suppress the AOA community [88], although this antagonistic relationship may be alleviated in N-rich soil [90]. Because AOA play an important role in the first step of nitrification [85, 91], AMF effects on AOA abundance and activity have implications for terrestrial N cycling and N2O emissions.

### Interactions between AMF and decomposers that can potentially degrade fungal or plant biomass

AMF hyphae turnover quickly (5-6 days, [92]) and cycling of this necromass represents a potentially rapid flow of nutrients and photosynthate C into a large volume of the soil system [43, 93]. Our study’s highly ^13^C-enriched MAGs have the genomic potential to degrade AMF fungal biomass. All of the enriched taxa contain GH109 genes, whose primary reported activity is α-N-acetylgalactosaminidase (additional functions may be as yet unknown). Galactose is a component of polysaccharides attached to the surface of AMF hyphae [79], and a lectin specific to D-galactose or N-acetyl-D-galactosamine glycoproteins was able to bind strongly to protein extracted from AMF [94]. In particular, Fibrobacterota mCT95 has 11 putative chitinase genes (GH18, GH19) and appears to be deriving most of its C from the AMF (mCT95 has the same APE-^13^C as the AMF), and thus could be performing a chitinolytic function in this soil food web.

All the highly enriched MAGs and many of the low- to medium-enriched MAGs have enzymatic potential to degrade components of plant biomass. The MAGs from Fibrobacterota and Verrucomicrobiota family Opitutaceae have isolated relatives that are thought to be involved in decomposition. Fibrobacterota include cellulose degrading bacteria found in mammal rumens [95], termite guts [96], anaerobic cellulose reactors [97], and rice paddy soil [98]. The Verrucomicrobiota family Opitutaceae contains isolates derived from rice patties and insect guts [99–102]. In our previous study decomposition gene expression in the rhizosphere and detritusphere using metatranscriptomics, we found that Fibrobacterota and Opitutaceae were two of the three groups that exhibited the highest decomposition gene expression when both root exudates and detritus were available [103]. Further examination would be required to determine if the MAGs from this study exhibit similar synergistic behavior in the AMF hyphosphere, and whether this stimulates the decomposition of plant residues.

### Methodological considerations: SIP DNA recovery

The amount of DNA recovered in each SIP density fraction can be a limiting factor for library construction, especially in the highest and lowest density fractions where only a small portion of total DNA is captured. In our research, we typically aim for 100 ng DNA per fraction, so that 20 ng may be reserved for 3 analyses: metagenomic sequencing, 16S rRNA gene and ITS amplicon libraries, and qPCR assays. To reliably achieve ∼100ng DNA in the majority of the fractions collected, we typically recommend loading 3-5 µg of DNA, because not all DNA added to the density centrifuge tube is recovered after fraction collection and desalting. Prior to adding non-ionic detergents, we noticed that DNA recovery appeared to be correlated with the initial amount loaded and hypothesized that DNA was being lost by an adsorption mechanism. Adsorption of DNA to polypropylene tube walls can potentially lead to substantial sample loss, especially when DNA is in a high ionic strength solution [48], such as CsCl gradient buffer. Our tests indicate that adding a low concentration of Tween-20 (0.0001%) leads to a near doubling of DNA yield. Adding Tween-20 may be particularly critical when limited DNA is available, such as from small samples or low biomass environments. Using this method, samples with 1 µg DNA and below can be more reliably processed and analyzed using metagenomic sequencing technologies that have low DNA input requirements. Here, we successfully analyzed 350 ng DNA per hyphosphere DNA sample with a 42% DNA recovery, which was suitable for metagenomic sequencing of the fractions containing DNA.

### HT-SIP optimization

Automating the SIP process significantly decreased the required operator hours (Table 1) while simultaneously improving reproducibility and sample recovery. Traditionally, after manually fractionating a density gradient, an additional 1-2 hours per sample is required to desalt gradient fractions (nucleic acid purification). Thus, in many research groups, a maximum of 6-8 samples may be processed per week, a grueling prospect if this pace is kept up week after week. HT-SIP makes it possible to routinely process 16 samples on a weekly basis, since the overall time to process a group of samples is decreased by over half, and the laborious “hands-on” tasks are significantly decreased by one-sixth. These time savings also translate into substantial labor cost savings, which over time can offset the initial cost of purchasing robotic instruments.

In the process of assessing different time saving methods, we attempted different techniques that were not adopted as part of the final pipeline, but these experiences may benefit others when developing their own pipelines. In addition to PEG precipitation, we attempted magbead cleanups, which are more time efficient than PEG if multiple magnetic plates are run in parallel; however in our hands, we found that PEG precipitations had higher yields. We also assessed if DNA intercalators could be used to minimize density differences associated with differences in GC content, with the goal of minimizing the need for separate 12C controls. Actinomycin D has been shown to reduce the native buoyant density of DNA with greater effect in GC rich DNA [104], thus theoretically reducing 90% of natural density differences. We found that the high concentrations needed to reduce the density of GC rich DNA also reduced the quality of the DNA density distribution and the overall DNA recovery. While we did not pursue further use of intercalators, with some optimization this could be a viable approach. These two protocols are available in the Supplemental Methods.

## CONCLUSIONS

Increasing the throughput of SIP is needed to promote well-replicated ecological scale studies to determine the ecophysiology of uncultivated organisms in complex environments. Here, we demonstrated an automation approach to expedite the most tedious tasks for SIP—fractionation, cleanup, quantification—that can increase the throughput and decrease the variability of SIP, with DNA recovery that is comparable to manual SIP processing. Decreasing the hands-on labor needed to run SIP samples inherently makes high-risk samples more feasible, such as the AMF hyphosphere samples we analyzed, where we had limited soil volume, low bulk atom percent enrichment and minimal separation based on total DNA density curves. The highly ^13^C-enriched hyphosphere MAGs identified in this study highlight the potential for trophic interactions in this zone, which includes predation, decomposition of fungal or plant biomass, and ammonia oxidation. In combination with other ‘-omics technologies, such as metatranscriptomics or proteomics, these MAGs will provide an important genomic resource for future experiments exploring interactions between AMF and their native microbiome.

## ACKNOWLEDGEMENTS

We thank Steve Kubala for assistance programming the robotic methods, Craig See for assistance with manual refractometry, G. Mike Allen for SIP technical assistance, and the JGI IMG and metagenomics teams for assistance with data processing (Neha Varghese, Alicia Clum, Marcel Huntemann, Tatiparthi Reddy, Supratim Mukherjee). Development of the HT-SIP pipeline was sponsored by the Joint Genome Institute through an Emerging Technologies Opportunities Program award (DOI: 10.46936/10.25585/60008401) to J.P.R. Testing of experimental samples on the LLNL HT-SIP pipeline was supported by the U.S. Department of Energy Office of Science, Office of Biological and Environmental Research (BER) Genomic Science Program (GSP) ‘Microbes Persist’ Scientific Focus Area award SWC1632 to J.P.R. The ^13^CO_2_ plant-AMF experiment was supported by DOE BER GSP awards DE-SC0016247 and DE-SC0020163 to M.K.F, and hyphosphere-SIP metagenomics sequencing and analysis was supported by DOE BER Early Career award SCW1711 to E.E.N. Work at Lawrence Livermore National Laboratory was conducted under the auspices of the U.S. DOE under Contract DE-AC52-07NA27344. The work conducted by the U.S. Department of Energy Joint Genome Institute (https://ror.org/04xm1d337), a DOE Office of Science User Facility, is supported by the Office of Science of the U.S. Department of Energy operated under Contract No. DE-AC02-05CH11231.

