## Supplemental Figure S1 for "HT-SIP: A semi-automated Stable Isotope Probing pipeline identifies interactions in the hyphosphere of arbuscular mycorrhizal fungi"

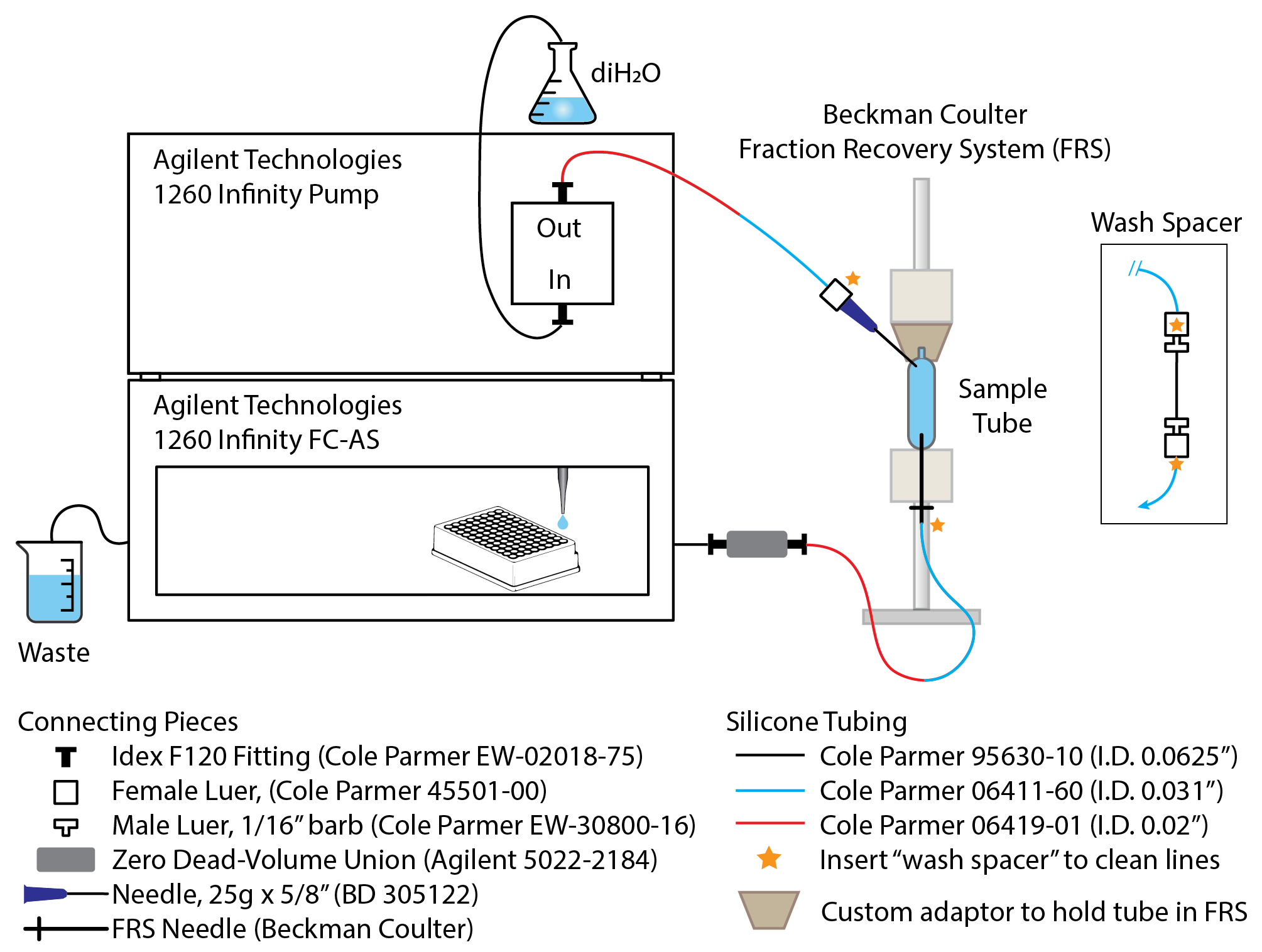

**Supplemental Figure 1. Automated fractionation system design, key parts and assembly.** We automated density fractionation by connecting an Agilent Technologies 1260 Infinity Fraction Collector to a Beckman Coulter Fraction Recovery System (FRS); part numbers are provided for the tubing and connectors required to build this system. The Agilent Infinity pump displaces the gradient buffer with a displacement fluid (e.g., sterile deionized water), and the gradient buffer fractions are collected in a 96-well plate. We attached a custom adaptor to the FRS to hold the gradient tubes in place, since we do not flow the displacement fluid through the FRS directly but instead puncture the top of the tube with a needle and flow the displacement fluid directly into the SIP sample tube. To create the custom adaptor, we use a rubber stopper or cork with a hole on the bottom hollowed out to hold the tube neck, and a piece of plastic (e.g., syringe bore) inserted into the top of the stopper that slots into the FRS gasket; we then hold the tube in place by compressing the FRS upper mount and this adaptor on top of the tube. To clean the tubing after fractionating a sample, the tubing is detached from the needles at the yellow stars and then attached to the wash spacer (attaches to yellow stars on inset diagram); the tubing is then flushed with water and the wash water is collected as waste. Mineral oil can also be used as a displacement fluid.
